# Bioprospecting Native Oleaginous Microalgae for Wastewater Nutrient Remediation and Lipid Production

**DOI:** 10.1101/2025.09.10.675442

**Authors:** Bobby Edwards, Daris P Simon, Ashish Pathak, Devin Alvarez, Ashvini Chauhan

## Abstract

Subtropical climate in Florida offers a unique opportunity for sustainable biofuel production using native microalgae cultivated in untreated wastewater. This study bioprospected oleaginous microalgal consortia from a wastewater holding tank at the Thomas P. Smith Water Reclamation Facility in Tallahassee, Florida, aiming to identify strains capable of both nutrient remediation and lipid accumulation. Using Fluorescence Activated Cell Sorting (FACS), chlorophyll-containing cells were isolated and cultured on BG-11 media. Shotgun metagenomics revealed that the most robust consortia- labeled C3, C4, and C9, were dominated by *Chlamydomonas*, *Acutodesmus*, and *Volvox* spp., alongside diverse bacterial, fungal, and archaeal communities. Functional gene analysis indicated active pathways for photosynthesis, lipid biosynthesis, and nutrient assimilation. In microcosm experiments, these consortia achieved up to 100% ammonia, 95% phosphorus, and 89% nitrate removal, outperforming control treatments. Lipid screening confirmed significant accumulation, with consortium C9 showing the highest yield. These findings underscore the potential of native microalgal consortia for integrated wastewater treatment and biofuel production, advancing circular bioeconomy strategies for subtropical regions.

## Introduction

The increasing research for biofuel development is a global collaborative effort to combat climate change, respond to higher energy consumption, and secure energy supply to meet the demand of the increasing world population which is projected to rise above 10 billion by 2050 (Neste Corporation, 2016). Biomass energy lies at the heart of renewable energy needs due to the global distribution of biomass, making the world demand for biomass energy (biofuel) grow by 41 to 53 billion liters over 2021 to 2026 (US Department of Energy, DOE, 2025).

Microalgae is one of the most promising sources of biofuels because it grows rapidly, yields large amounts of lipids, and can also remove pollutants while thriving in wastewater nutrients (Kozyatnyk, 2025). Microalgae as a feedstock for the renewable bioenergy production has the potential to meet the global biofuel demand as the lipid yield of some microalgae strains can exceed that of most productive oil crops (Wang *et al*., 2024). According to Williams (2007), a conservative estimate between 30,000 to 50,000 liters of lipids per hectare/ year can be obtained from microalgae cultivation compared with 1300 to 2400 liters per hectare/ year for plants with a high oil yield such as jatropha and palm oil, and the cost of biofuel production from microalgae is much cheaper than that from other biofuel sources (Williams, 2007).

The recent assessment of the potential benefits of microalgae has undoubtedly boasted the research focus on this subject, with many studies looking into the factors that could improve the biomass and lipid yield in controlled photobioreactors, thus reducing the risks associated with largescale outdoor cultivations (Wang *et al*., 2024). Several factors that influence the yield of biomass and lipids from microalgae cultivation have been largely investigated. In a recent study, Gopalakrishnan *et al*., (2024) applied response surface methodology (RSM) experimental design and reported that biomass yield from microalgae could increase up to 10^2^ cells/ml at the optimum conditions of CO_2_ level, light intensity, harvest time, and co-culture ratio of bacteria and microalgae, and the lipid yield could increase up to 10^8^ at the same optimum condition (Gopalakrishnan *et al*., 2024). Co-culture of bacteria and microalgae, through mechanisms ranging from mutualism to antagonism, has been reported as an important factor for the functionality and productivity of microalgal biomass and lipid yields (Berthold *et al.,* 2019). Dao *et al*. (2018) identified that bacteria co-cultured with microalgae can secrete Indole Acetic Acid (IAA) which enhances algal growth, and microalgae in turn can selectively enhance the secretion of IAA. Also, climatic conditions, such as the light intensity are critical to the overall metabolic activities of microalgae (Gao *et al*., 2024; Cabanelas *et al*., 2017). Nevertheless, the existing technologies seem not to have addressed the need to improve the economic viability of microalgae-based biofuel (Okoro *et al*., 2019).

Nutrients such as Nitrogen and Phosphorus in water are essential for plant growth but in excess, they can lead to significant problems, among such is accelerated eutrophication (Singleton, et al., 2022). Eutrophication occurs when a disproportionate amount of phosphorus and nitrogen enters the environment, mainly through animal fecal material, wastewater and sewage sludge, resulting in algal blooms causing a decline of aquatic life due to hypoxia and suffocation (Wright, 2022; Pinelli et al., 2022). Commonly, excess nutrients in wastewater are treated by chemical precipitation (Bunce et al., 2018). However, the Enhanced Biological Phosphorus Removal (EBPR) approach is gaining momentum due to the relatively low capital and operational costs and friendliness to the environment (Marques et al., 2017). EBPR requires the enrichment of wastewater with Polyphosphate Accumulating Organisms (PAOs), notable among whom are *Candidatus Accumulibacter Phosphatis*, *Candidatus Halomonas Phosphatis*, and *Pseudomonas putida* spp. (Zheng *et al*., 2022). An emerging aspect of EBPR often referred to as Photo-Enhanced Biological Phosphorus Removal (PEBPR) combined PAOs with algal species which relies on light/dark cycle to create the aerobic/anaerobic conditions that are needed to effectively remove excess nutrients in wastewater (Plouviez & Brown, 2024). However, a better understanding of the wastewater microbial communities, especially the microalgal-bacterial consortia that will effectively reduce the wastewater nutrient load will significantly reduce operation costs of the wastewater treatment plants and reduce eutrophication and its associated environmental challenges.

On the other hand, most of the previous studies on microalgae for biofuel production have relied on axenic or isolated monocultures, paying less attention to the potential of microalgae-microbial consortia (Faried et al., 2023; Ren et al., 2019; Dao et al., 2018). While these approaches have proven effective in academic research, they have shown limited industrial viability (Laezza *et al*., 2022). The productivity of mono algal cultures remains constrained due to the limited opportunities of optimization, and the available strategies such as nutrient supplementation, contamination prevention, novel bioreactor systems and genetic engineering, are not often cost-effective for large-scale applications (Laezza et al., 2022; Huy et al., 2018). Co-cultivating microalgae with other microorganisms that are either naturally present in their growth media or that are added externally has the potential to encourage cell development and the synthesis of a variety of very valuable compounds (Acién Fernández et al., 2018). The early 1950s saw the first mention of the symbiosis between algae and other microorganisms (i.e., bacteria) as a means of enhancing the oxygen (O_2_) supply to oxidation ponds in wastewater treatment plants (Oswald et al., 1953). Microalgae and other microbes have evolved a symbiotic relationship that includes all relationships known to exist in nature, including mutualism, commensalism, and parasitism (Ramanan et al., 2016; Su *et al*., 2022). The mechanism of the interactions among the communities in the consortia may be quite complex, but it is expected that the natural associations would enhance the algal metabolic activities, including the biofuel yields (Dao *et al*., 2018). It is however imperative to isolate high lipid producing microorganisms by bioprospecting different environmental niches and then evaluate their taxonomic and genomic traits prior to their successful application of wastewater remediation coupled with biofuels production.

Therefore, the objectives of this study are: (1) to isolate high lipid-accumulating algal consortia from the Thomas P. Smith Water Reclamation Facility (TPSWRF), located in Tallahassee, Florida and (2) to characterize the isolated strains to better understand their taxonomic lineages and gene potential and 3) to evaluate their ability to remediate wastewater such that a circular bioeconomic process that is environmentally sustainable can be developed for sub-tropical southern states such as Florida in the United States.

## Materials and Methods

### Wastewater Site Description and Sample Collection

Water samples were collected from the Thomas P. Smith Water Reclamation Facility (TPSWRF) in Tallahassee Florida. The facility is designed to receive municipal sewage from homes and businesses within Tallahassee Urban Service Area through the sanitary sewage collection system (Deltac & Meuser, 2017). The treatment process at the TPSWRF consists of preliminary treatment, primary clarification, biological treatment, secondary clarification, tertiary treatment and disinfection. The first two steps are meant to remove debris, sand, grease, oil and other solid particles before the biological treatment, which basically enhances microbial growth for the decomposition of the organic matters. Products from the biological treatment is stored in a pond (secondary clarifier) to separate solids from liquids before the chemical treatment and disinfection (Pfeffer, 2025).

At the holding pond of the secondary clarifier of the TPSWRF facility (Supplementary Figure S1), we collected three wastewater samples (3L each) using sterile plastic containers. The samples were tightly sealed and taken at room temperature to the Environmental Biotechnology Laboratory, School of Environment, Florida A&M University for further analysis.

### Cell Sorting Using FACS

The three wastewater samples S1, S2, S3, were filtered using 50μm mesh and then 80 ml of each sample was inoculated into 100 ml BG-11 growth medium and incubated at 25 ±2°C, for microbial enrichment with light cycle 16/8 hours on/off set at 1200 Lux for 7 days. This enrichment step is necessary to ensure that the fast-growing species are eventually selected over the slow-growing species (Pereira *et al*., 2011). We then applied a fluorochrome-BODIPY 505/515 (4,4-Difluoro-1,3,5,7-Tetramethyl-4-Bora-3a,4a-Diaza-s-Indacene), a high oil/water coefficient fluorescent dye developed to effectively screen lipid accumulation in microalgae, and can maintain its fluorescence longer than 30 min at the optimum concentration of 0.067 µg/ml (Govender *et al*., 2012). BODIPY 505/515 was dissolved in 0.2% dimethyl sulfoxide (DMSO) to a stock concentration of 1 mM and was added to stain the enriched cells at a final concentration of 1µM. The stained samples were then mixed gently and incubated in the dark at 25°C for 5 min before sorting in the Fluorescence Activated Cell Sorting (FACS) system. The FACS analysis was conducted with the four laser 10-color FACSCanto instrument at the Flow Cytometry Lab, College of Medicine, Florida State University, following the operational procedures (https://med.fsu.edu/flowCytometryLab/home).

Briefly, the BODIPY stained samples were loaded onto the FACS platform, and events logged for 5 min. To provide visual data, events were plotted by the side scattering angles against the fluorescence intensity; this data was visually inspected, and gates P2, P3, P4, P5 and P6 were generated on the resolved bands. For the first sample (S1), due to the events with higher relative fluorescence unit (RFU), gate P3 was sorted onto 96-well microtiter plates filled with solid BG-11 at a rate of 10 events per well based on the high event concentration. The FACS platform was washed with a 10% bleach solution after sorting S1, followed by sterile Milli-Q water, and then S2 and S3 were sorted like the S1 sample. For samples S2 and S3, Gate P2 and P6 were sorted into 96-well microtiter plates filled with solid BG-11, respectively. The sorting strategies for untreated wastewater samples S1, S2, and S3 are demonstrated in Supplementary Figures S2 to S4.

After sorting the samples, all the 96 well plates were incubated at room temperature, cultures were checked on a weekly basis, and signs of growth and contamination were visually monitored. Actively growing cultures were sub-cultured onto BG11 solid media under 50W glow bulbs. Several hundred isolates were recovered from the FACS experiment, and these were further narrowed to ten distinct samples that showed notably faster biomass development under the treatment conditions.

For further analysis, the ten best grown consortia from FACS were cultured and maintained in lab-scale bubble flasks (250 ml suspension volume each in BG-11 media) under continuous light, 14 hours day light (400 µmol photons m^−^² s^−1^) and 10 hours dark period, at 25±4 °C as the best practice (Rayati et al., 2020). The pH was measured regularly using the ST-3100 pH Meter (OHAUS Corporation, Parsippany, NJ, USA), and the biomass concentration was determined by measuring the optical density of samples at wavelength 750 nm (OD_750_) using an ultraviolet-visible range of spectrophotometer (SpectraMax M5, Molecular Devices, Sunnyvale, CA). These cultures were then subjected to lipid screening to find the suitable consortium for further analysis.

### Screening of the Algal-microbial consortia for Lipid Production

Nine consortial isolates obtained from the FACS analysis were subject to Sulfo-Phospho-Vanillin (SPV) screening for lipid accumulation (Mishra *et al*., 2014). Briefly, 10 ml of the algal cultures were centrifuged at 3000 rpm for 10 min. The supernatant was discarded and the residue dried in oven at 60°C. 3 mg of dried algae was scraped off and put in a pre-labeled 15ml centrifuge tube, 100 µL of sterile water was added to the tubes, followed by 2 mL sulfuric acid. The tubes were heated at 100°C for 10 minutes in a water bath (Buchi Waterbath B-481) and incubated in an ice bath for 5 minutes. After the incubation, 5 mL SPV was added to the tubes and kept on a shaking incubator at 37°C for 15 minutes at 200 rpm. The absorbance was measured in triplicates at 530nm by micro-volume spectrophotometer (Eon et al., V.T, USA). The lipid yield was calculated using a triolein standard calibration (R^2^ =0.97). Consortia with over 50% (dry weight) lipid yield were selected for further analysis.

### Sequencing and Bioinformatics Analysis of the Isolated Strains

Genomic DNA from actively growing strains labeled as consortia 3, 4 and 9 and a mix of all consortia was extracted using the DNeasy PowerLyzer Microbial Kit, following the manufacturer’s directions (Qiagen, Germantown, MD, https://www.qiagen.com/us/products/). Sequence libraries for shotgun metagenomics were prepared utilizing the Illumina Nextera XT kit, according to the manufacturer’s instructions (Illumina, San Diego, CA). Sequencing was performed on an Illumina NextSeq500 instrument employing a mid-output kit with 2×150 paired-end sequencing. Raw reads were mapped to the NCBI non-redundant protein database for taxonomic profiling using DIAMOND (Buchfink *et al*., 2015). Taxonomic summaries per read were obtained using MEGAN’s Least Common Ancestor algorithm (Huson *et al*., 2007) and then summarized across all reads to create counts per taxon. Functional profiling at subsystem levels 1 and 3 was performed using SUPER-FOCUS (Silva *et al*., 2016), and raw counts were normalized to counts per million (CPM) units for relative abundance estimates. Stacked bar plots were generated on the relative abundance estimates from each sample at the phylum and genus levels separately for bacteria, archaea, and fungi. For the pure strains, sequencing for the stains identification was carried out using the Illumina MiseqV3 at the Carver Biotechnology Center (CBC) University of Illinois, USA. The DNA was extracted using the Bio-Rad DNA extraction kits and the genetic primers for the amplicon analysis were cyano-specific CYA106f and CYA781R. Raw reads were mapped to the NCBI non-redundant protein database for taxonomic profiling and then summarized across all reads were used to create taxon pie chart.

### Nutrient Depletion from Wastewater using the isolated consortia

The isolated pure strains were cultured and maintained in lab-scale bubble flasks (250 ml suspension volume each in BG-11 media) under continuous light, 14 hours day light (400 µmol photons m^−^² s^−1^) and 10 hours dark period, at 25±4 °C for 10 days as the best practice (Rayati et al., 2020). The strains were centrifuged at 5000 rpm for 10 min, a safe condition centrifugal without much cell disruption (Xu *et al*., 2015). The cell biomass was collected, and wet weight was taken. Wastewater samples were collected from the Thomas P. Smith Water Reclamation Facility (TPSWRF) in Tallahassee Florida, which were spiked with algal consortia and measured for depletion in nitrate and phosphate for 12 days. Briefly, 200 ml of triplicate wastewater samples were autoclaved and added into plastic buckets. Then 2 g wet weight of the algal cells were inoculated into the wastewater samples. Samples were then collected daily for 12 days and measured for total phosphate (TP) and total nitrate (TN) analysis.

### Phosphate, Ammonia and Nitrate Analyses

Total phosphorus in the wastewater samples was analyzed using the Inductively Coupled Plasma Optical Emission Spectrometry (ICP-OES) at the core lab facility, School of the Environment, Florida A&M University. Samples were first digested by adding 1 ml of concentrated H_2_SO_4_ and 0.5 g potassium persulfate (K_2_S_2_O_8_) followed by heating in a water bath at 100°C for 1 hour, followed by filtration using 0.45 µm membrane. The filtrate was then injected into the machine and the total phosphorous was measured at 213 nm against phosphorous ICP standard solutions. The phosphate concentration was calculated by multiplying the total phosphorus by 3.06.

Nitrate analysis was carried out colorimetrically using a UV spectrophotometer at 220 nm and 275 nm. The absorbance difference was calculated by subtracting the absorbance corresponding to organic matter interference at 275 nm from the absorbance at 220nm (Karlsson *et al.,* 1995). A calibration standard was prepared using potassium nitrate (KNO_3_) standard solution. Wastewater dissolved ammonia was analyzed using a handheld photometer (Lumiso Ammonia Photometer, https://www.waterindustryjournal.co.uk/).

### Metagenomic Sequence Accession Numbers

The shotgun metagenomic sequence data obtained from this study are available from NCBI’s Sequence Read Archive/European Nucleotide Archive, accession under BioProject ID PRJNA1198942, Biosample numbers, and SAMN45875459, SAMN45875460, SAMN45875461 and SAMN45875462, respectively. A direct link to the data is as follows: https://submit.ncbi.nlm.nih.gov/subs/sra/SUB14932812/overview.

## Results and Discussion

### Isolation of Microalgae from Wastewater Samples

Using Fluorescence Activated Cell Sorting (FACS) on enriched wastewater samples, we were able to isolate hundreds of green pigmented potentially microalgal strains, as shown in supplemental Figures 1-5. We believe that this approach is more applicable for the quick isolation of environmental lipid producing strains relative to the traditional culture-based laboratory approach, which is labor intensive and generally takes much longer time (Laezza *et al*., 2022). We expect that other researchers will also apply the FACS technique leading to a high throughput isolation system yielding high lipid producing isolates.

Sulfo-Phospho-Vanillin (SPV) method was then applied to screen the isolated nine strains using triolein standard calibration. Figure 1 shows that consortium C9 had the highest lipid accumulation, followed by C3 and C4, with lipid yields of 91, 86, and 75 µg/3mg, representing 3.1, 2.9, and 2.5% (w/w), respectively. Other consortia had lipid yield less than 2.0%, hence consortia C9, C3 and C4 were selected for further analysis. The lipid contents of these consortia were much below the average for some notable high lipid-producing microalgae such as *Nannochloropsis*, *Scenedesmus*, and *Schizochytrium* whose lipid yield ranges from 37–60, 30–50, and 50–77 wt.%, respectively (Shivakumar, *et al*., 2024). Therefore, growth optimization studies are suggested to optimize the lipid productivity from the newly isolated consortia.

**Figure 1.**
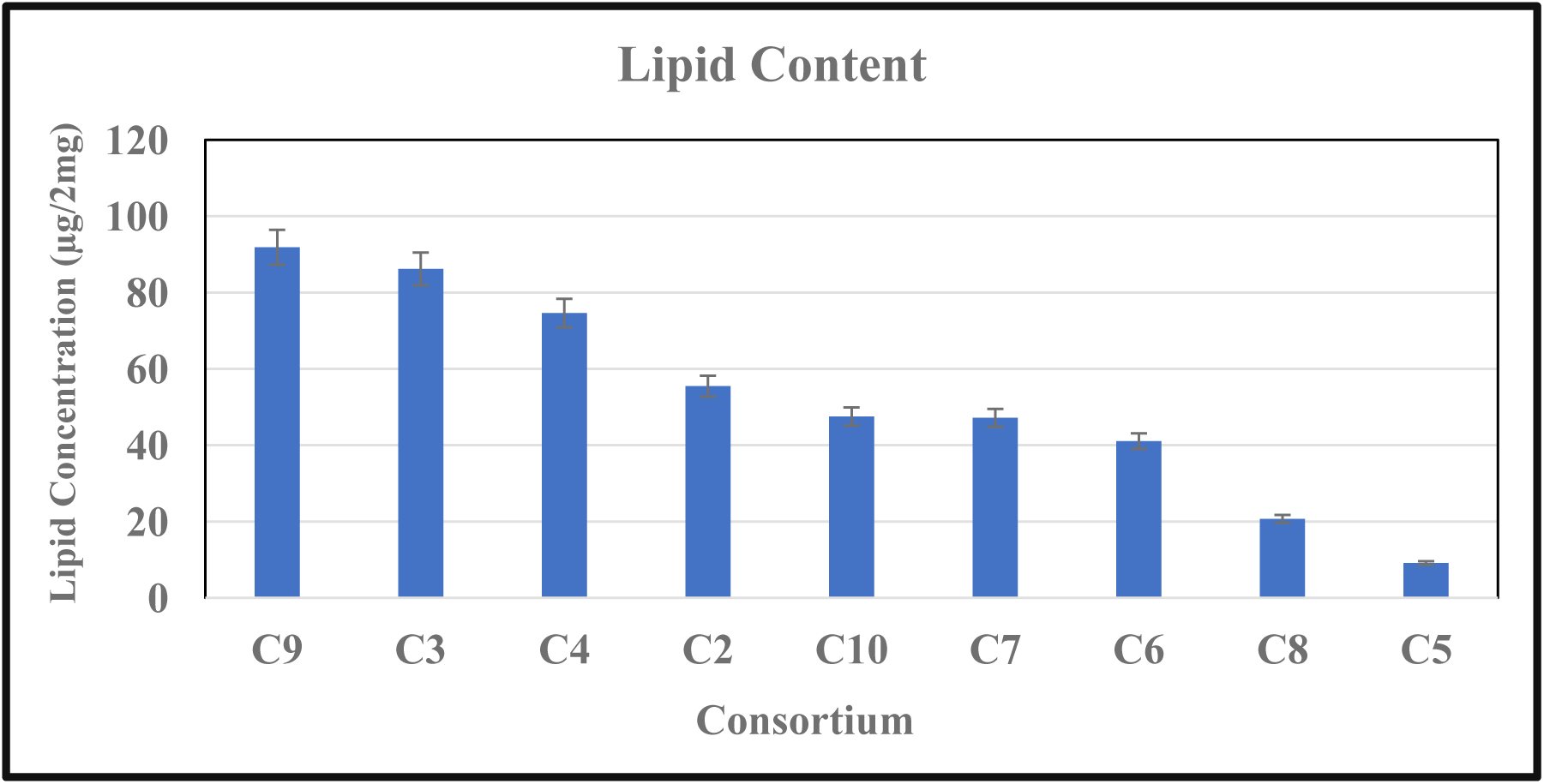
The lipid yield of the microalgal consortia isolated from the TP Smith Wastewater Reclamation Facility, Tallahassee, FL.

### Identification of the Wastewater Consortium

Shotgun metagenomics was performed to evaluate the algal consortium taxonomy and gene functions. Once the abundance files from each consortia was obtained, it was analyzed using the microbiomeanalyst pipeline (Dhariwal et al., 2017). Data were filtered for low count and low variance using default parameters, which resulted in the removal of 132 low abundance features based on prevalence and 126 low variance features based on Interquartile Range (IQR). A total of 1128 features remained for downstream processing after the data filtering step. Sequences were then rarefied to the minimum library size, and total sum scaling was performed to account for sample variability so that biologically meaningful comparisons could be drawn.

As shown in Figure 2, regardless of the sample, the algal consortia were dominated by *Chlamydomonas* spp., which form the core microbiome in these samples. Notably, *Chlamydomonas* has been found in diverse ecological niches ranging from fresh to marine environments and wastewater (Kong et al., 2010). Moreover, *Chlamydomonas* are known for their rigorous biodegradative abilities in the presence or absence of bacteria (Sun & Simsek, 2017). Another group working on phytoremediation of wastewater found that *Chlamydomonas* was a prime candidate for removing the nutrients and significantly decreasing the amount of nitrate and orthophosphate (Shaker et al., 2015).

**Figure 2.**
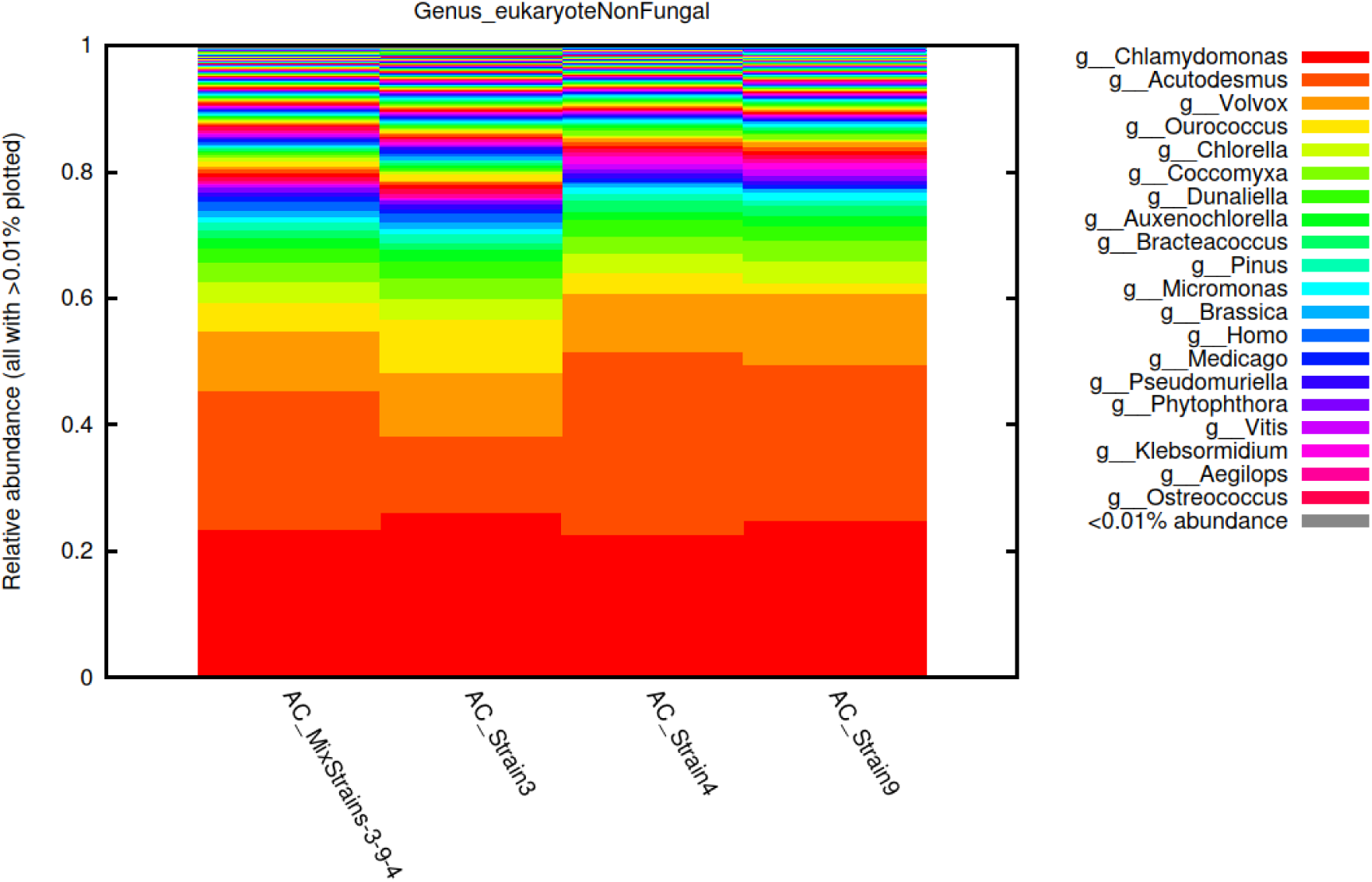
Shotgun metagenomics analysis showing the taxonomic composition of eukaryotic non-fungal communities in the oleaginous consortia isolated from the TP Smith Wastewater Reclamation Facility, Tallahassee, FL.

In addition to *Chlamydomonas*, our algal consortia was represented by at least 20% of *Acutodesmus* spp.-which has also been studied to reduce nutrients in wastewater from anaerobic digesters using mixtures of agro-industrial wastes (Koutra et al., 2017). The third genera in our consortia was represented at about 15% by *Volvox sp*., which has been identified to uptake nutrients in freshwater (Goldstein, (2015). With these diverse groups of algae, it can be hypothesized that they are working in tandem to uptake wastewater nutrients and produce lipid precursors for biodiesel production.

### Bacterial, Archaeal and Fungal Assemblages in the Consortia

Shotgun-based metagenomic analysis revealed that the microalgal consortium used in this study was mainly comprised of 5 phyla that were greater than 90% of the total relative abundance. Among these were *Pseudomonadota* (synonym “*Proteobacteria*”), which accounted for over 70% of the significant diversity, followed by *Actinomycetota* (or *Actinobacteria*), *Bacteroidota*, and *Bacillota* (data not shown). *Pseudomonadota* have been commonly found dominating bacteria phyla in microalgae consortia, possibly due to the carbon degradation ability of *Pseudomonadota*, requiring oxygen from the microalgae and supplying CO_2_ in a symbiotic relationship (Rodriguez *et al*., 2023). Similar studies by Numberger *et al*. (2019) in a wastewater treatment plant in Berlin, Germany, also showed these phyla to be predominant wastewater microorganisms. Furthermore, 16S identification of a wastewater microbial community in Shanxi, China, also showed the abundance of *Pseudomonadota* and *Actinomycetota,* with the members of the genus *Thiobacillus* and *Comamonas* dominating the overall bacterial population (Ma *et al*., 2015).

At the genus level, it was observed that *Pseudomonas*, *Brevundimoas*, *Blastomonas*, and *Rhodobacter* were the most prevalent genera in the consortia, consisting of over 40% of the group (Figure 3). It does appear that the bacterial community is diverse and has a mutualistic symbiotic relationship with the algae. Along the lines of our findings, Numberger *et al*. (2019) also concluded that *Pseudomonas* was a major environmental microorganism in wastewater treatment compared to the whole genomic community. Likewise, Pastore and Sforza (2018) identified a symbiotic relationship between *Chlorella protothecoides* and *Brevundimonas* in which nutrients were exchanged/converted for growth during water remediation. In addition, Lee *et al*. (2017) discovered *Blastomonas* to be a novel species related to wastewater treatment that also has high fatty acid content. Finally, *Rhodobacter* is known to have diverse metabolic activities based on the environment, and one pathway is bioremediation (Costa *et al*., 2017).

**Figure 3.**
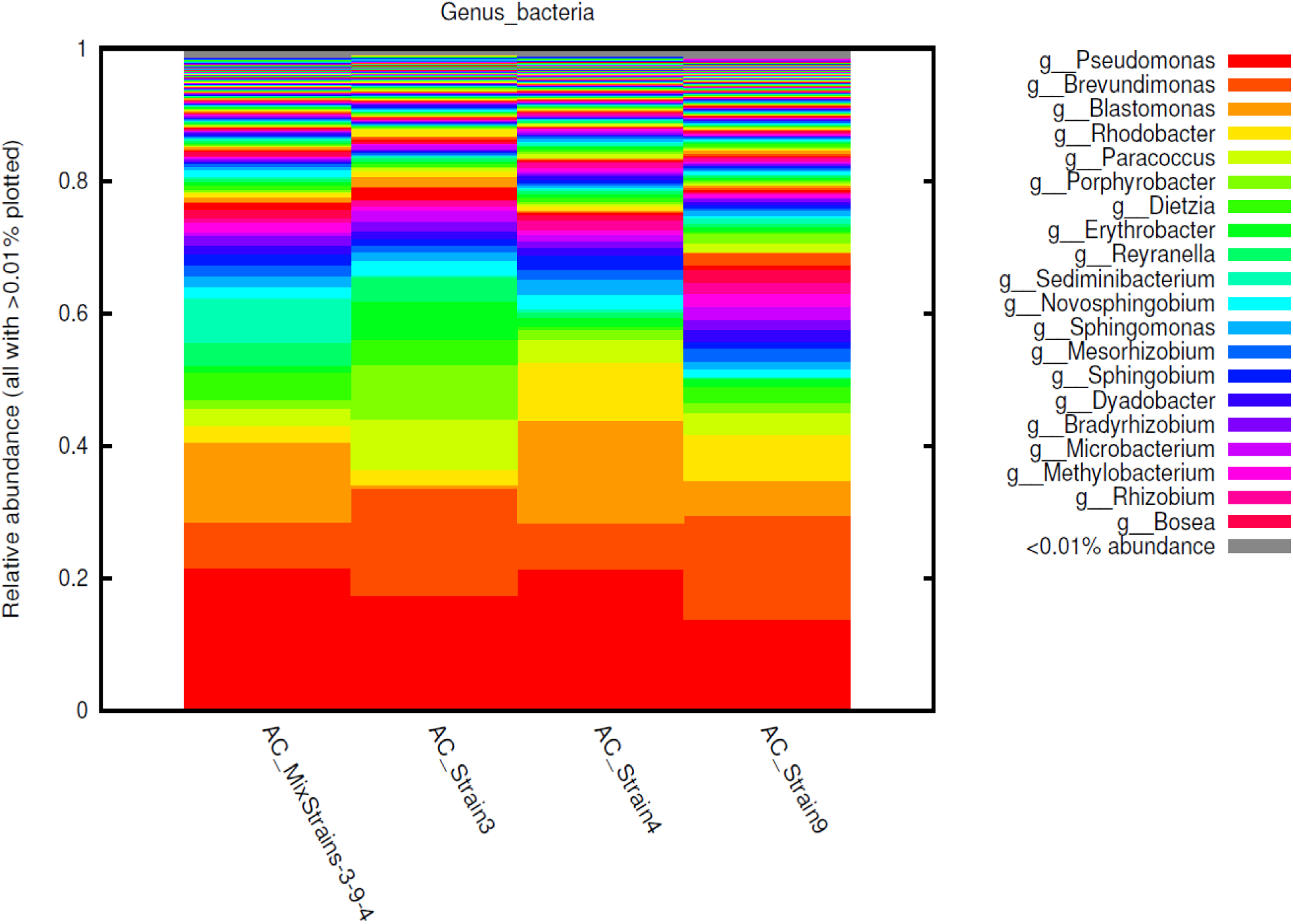
Shotgun metagenomics analysis showing the taxonomic composition of bacterial communities in the oleaginous consortia isolated from the TP Smith Wastewater Reclamation Facility, Tallahassee, FL.

Among the archeal members, phyla *Euryarchaeota*, *Thaumarchaeota*, and *Crenarchaeota* dominated the archaeal group in the consortia (data not shown). The percentage abundance of the phylum *Euryarchaeota* in C4 and C9 was over 80% and the phylum *Thaumarchaeota* was about 25% on the mixed consortia. However, Jabari *et al*. (2016) found that the phylum Euryarchaeota was only 8.9% of the microorganisms in wastewater and 70% less than the identified phyla in the consortia. Antwi *et al*. (2017) found that the phylum *Euryarchaeota* was the second most abundant group in wastewater at 22%, so *Euryarchaeota* is a consistent member of wastewater algal consortia. In addition, the phylum *Thaumarchaeota* was only identified to make up to 5.4% of the total community in municipal wastewater (Meerbergen et al., 2017). The phylum *Crenarchaeota* was low in the consortia because it has been previously discovered to have less tolerance to wastewater (Cortez-Lorenzo *et al.,* 2014; Gray *et al.,* 2002).

At the genus level, the archaeal members of the consortium community mainly consisted of *Methanocaldococcus* spp. (Figure 4), followed by *Halorubrum* and *Haloferax* spp. Notably, *Methanosarcina* spp. has been reported to enhance the generation of n-alkane-rich biofuels from microalgae (Yamane *et al.,* 2013). It is highly likely that *Methanosarcina* spp. play critical roles within the wastewater treatment, especially under anaerobic conditions (Tabatabaei et al., 2010). *Halorubrum* is an extremophile that grows in harsh environments such as Salt Lakes in Africa and the Antarctic ice (Karray et al., 2021; Salwan & Sharma, 2020), and such trait could explain why it overcome the harsh wastewater environment. Another microorganism that is known for bioremediation is *Haloferax* spp, which has been found with an anaerobic metabolism pathway for denitrification in various media (Torregrosa-Crespo et al.,2016). In all, the archaeal members of the consortia were already known for their crucial ecological roles in the community.

**Figure 4.**
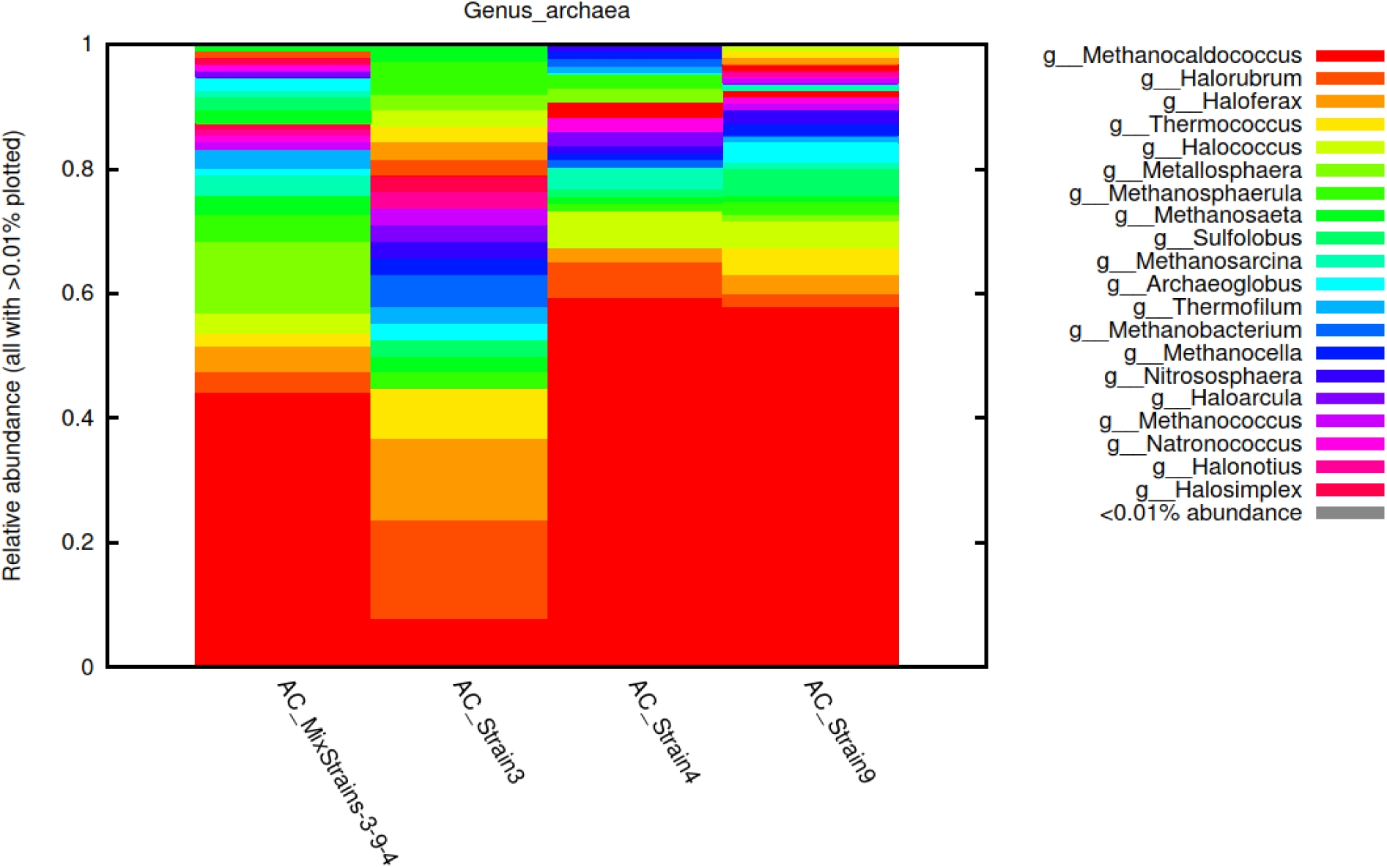
Shotgun metagenomics analysis showing the taxonomic composition of archaeal communities in the oleaginous consortia isolated from the TP Smith Wastewater Reclamation Facility, Tallahassee, FL.

Looking at the fungal community in the consortia, the *Ascomycota* and *Basidiomycota* phyla dominated over 97% of the total relative abundance of the fungal members (data not shown). Other members of the community were Glomeromycota and Chytridiomycota, *Cryptomycota* and *Blastocladiomycota*, being present in exceptionally low abundance in the consortia. There are comparable results in a South African study, where three wastewater treatment plants were sampled, and the fungal members of the consortia were dominated by *Ascomycota* and *Basidiomycota* (Assress *et al*., 2019). At the genus level, *Fusarium* spp dominated (Figure 5). *Fusarium*-*Chlorella* consortium has treatment of wastewater better than monocultures (Zhang *et al*., 2023).

**Figure 5.**
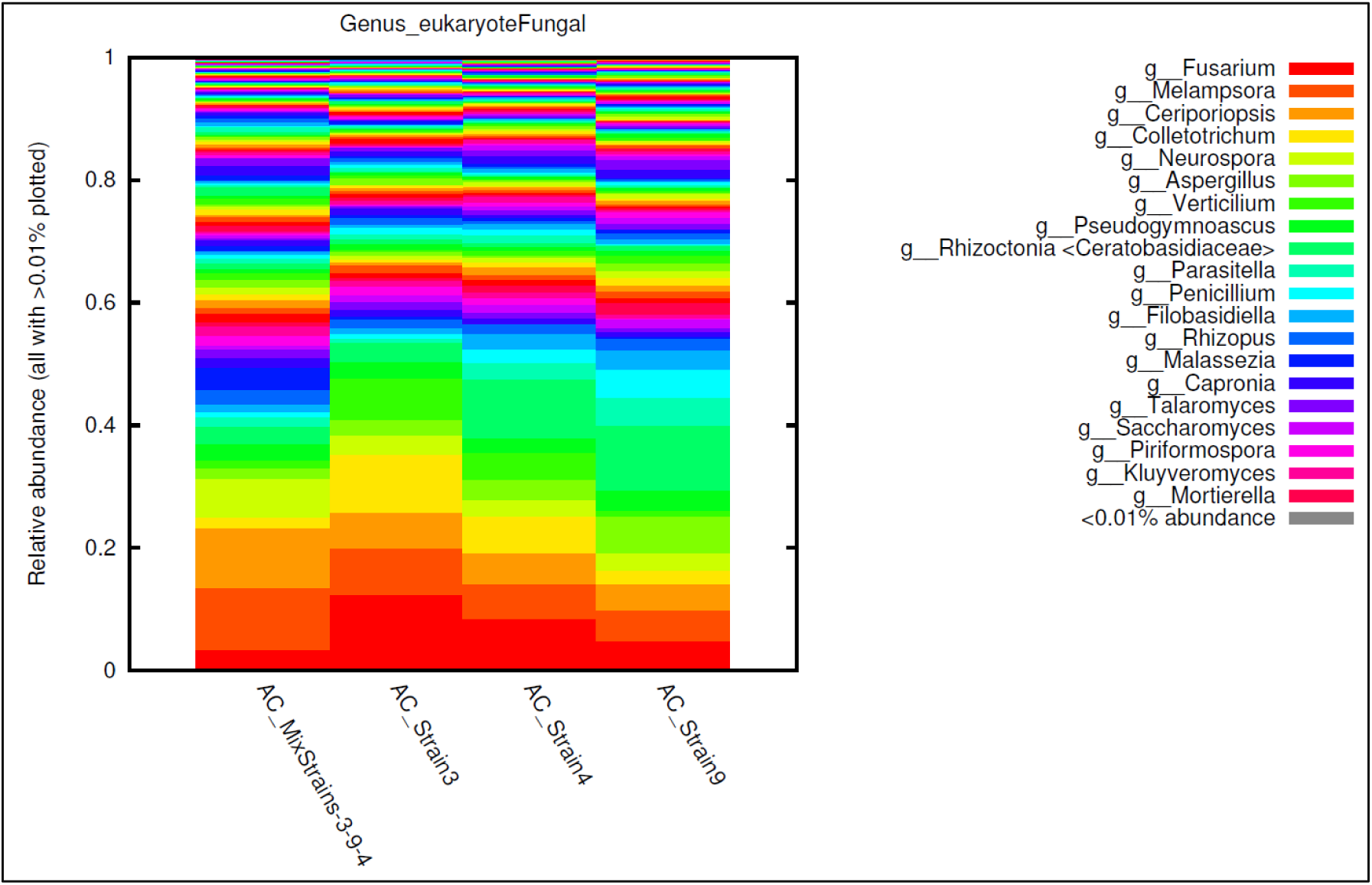
Shotgun metagenomics analysis showing the taxonomic composition of fungal communities in the oleaginous consortia isolated from the TP Smith Wastewater Reclamation Facility, Tallahassee, FL.

### Gene Functional Analysis in the Consortia

We also investigated the gene functions of algal, bacterial, archaeal, and fungal communities in the consortium. As shown in Figure 6A, functional metagenomics at subsystem level 1 indicated that genes performing functions related to carbohydrate metabolism, photosynthesis, protein metabolism, amino acids, and derivates, and respiration were the most abundant functions being performed in the consortium. Although at lower levels, functions related to stress response, membrane transport, fatty acids, lipids and isoprenoids, and metabolism of aromatic compounds were also observed in the consortium.

**Figure 6.**
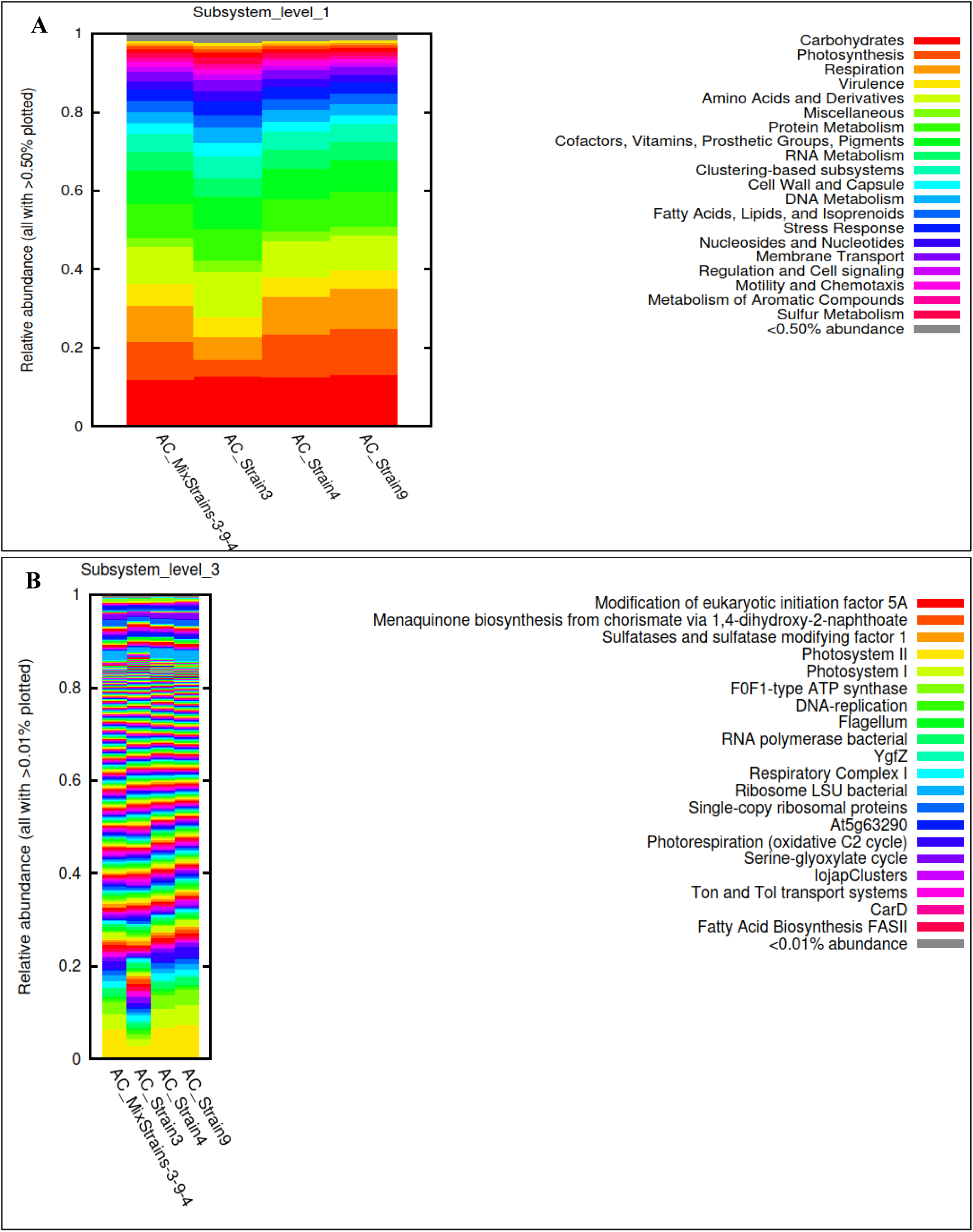
Shotgun metagenomics analysis showing the suite of genes present in the oleaginous consortia isolated from the TP Smith Wastewater Reclamation Facility, Tallahassee, FL. Shown are A, the functional genes at subsystem level 1 with genes performing functions related with carbohydrate metabolism, photosynthesis, respiration, virulence, amino acids and derivates and respiration as the most abundant functions being performed in the consortia; B, the gene functional analysis at subsystem level 3 which revealed gene classes known to facilitate the growth of algae, including the photosystem I and II, YgfZ proteins, photorespiration, oxidative C2 cycle, and fatty acids biosynthesis FASII in the consortia.

It was noteworthy that at subsystem level 3, several gene classes known to facilitate the growth of algae, including the photosystem I and II, YgfZ proteins, photorespiration (oxidative C2 cycle), and fatty acids biosynthesis FASII were observed in the consortia (Figure 6B). The high abundance of carbohydrate metabolic genes in microalgae are required for biofuel production and cell maintenance (Andreeva, et al., 2021).

### Wastewater Remediation by the Isolated Consortia

Consortia 3, 4 and 9 were evaluated for their ability to utilize wastewater nutrients, specifically phosphate, ammonia and nitrate. As shown in figures 7, 8 and 9, the phosphate, nitrate, and ammonia were effectively consumed by the consortial members over a 12-day period. For phosphorus (P), the experimental data demonstrated varying efficiencies of (P) removal from wastewater by three microbial consortia—Consortia 3, 4, and 9—compared to control, which did not contain any strain but just wastewater (Fig. 7). Consortium 9 exhibited the highest phosphorus utilization, showing a rapid and near-complete depletion of P over the course of the experiment. This suggests a robust metabolic capacity and strong adaptation to wastewater conditions. Consortium 4 showed moderate efficiency, with a steady decline in phosphorus levels, indicating effective but less aggressive uptake. In contrast, Consortium 3 displayed the lowest performance, with slower and less complete phosphorus removal. P in the control experiment also declined but the consortial treatments reduced P at a significantly higher rate. These findings highlight Consortium 9 as a promising candidate for bioremediation and phosphorus recovery applications.

**Figure 7.**
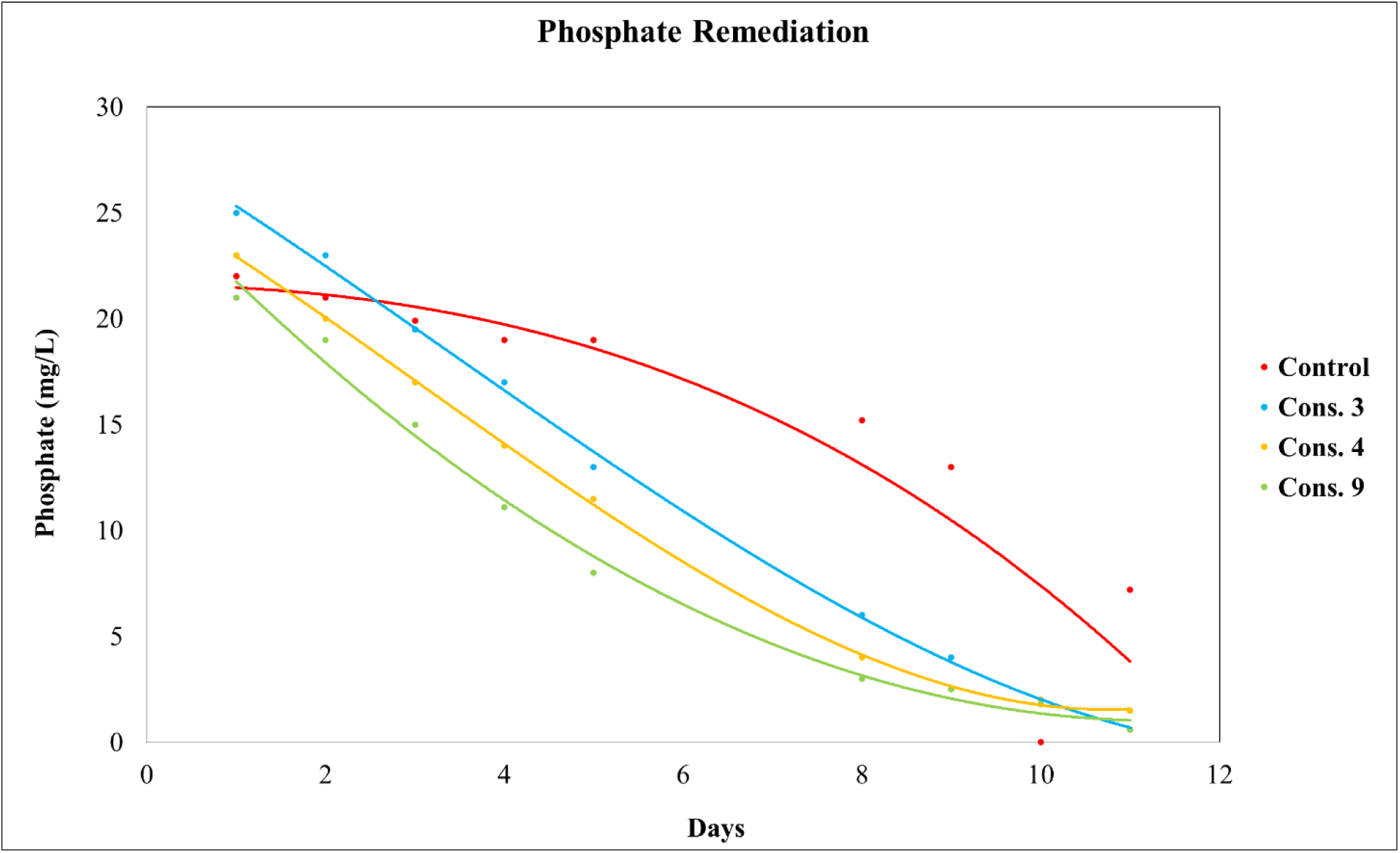
Phosphate removal by the oleaginous consortia isolated from the TP Smith Wastewater Reclamation Facility, Tallahassee, FL. The phosphorus content before and after treatment was analyzed using the Inductively Coupled Plasma Optical Emission Spectrometry (ICP-OES). A control treatment was included with only unautoclaved wastewater sample without addition of isolated consortial strains.

**Figure 8.**
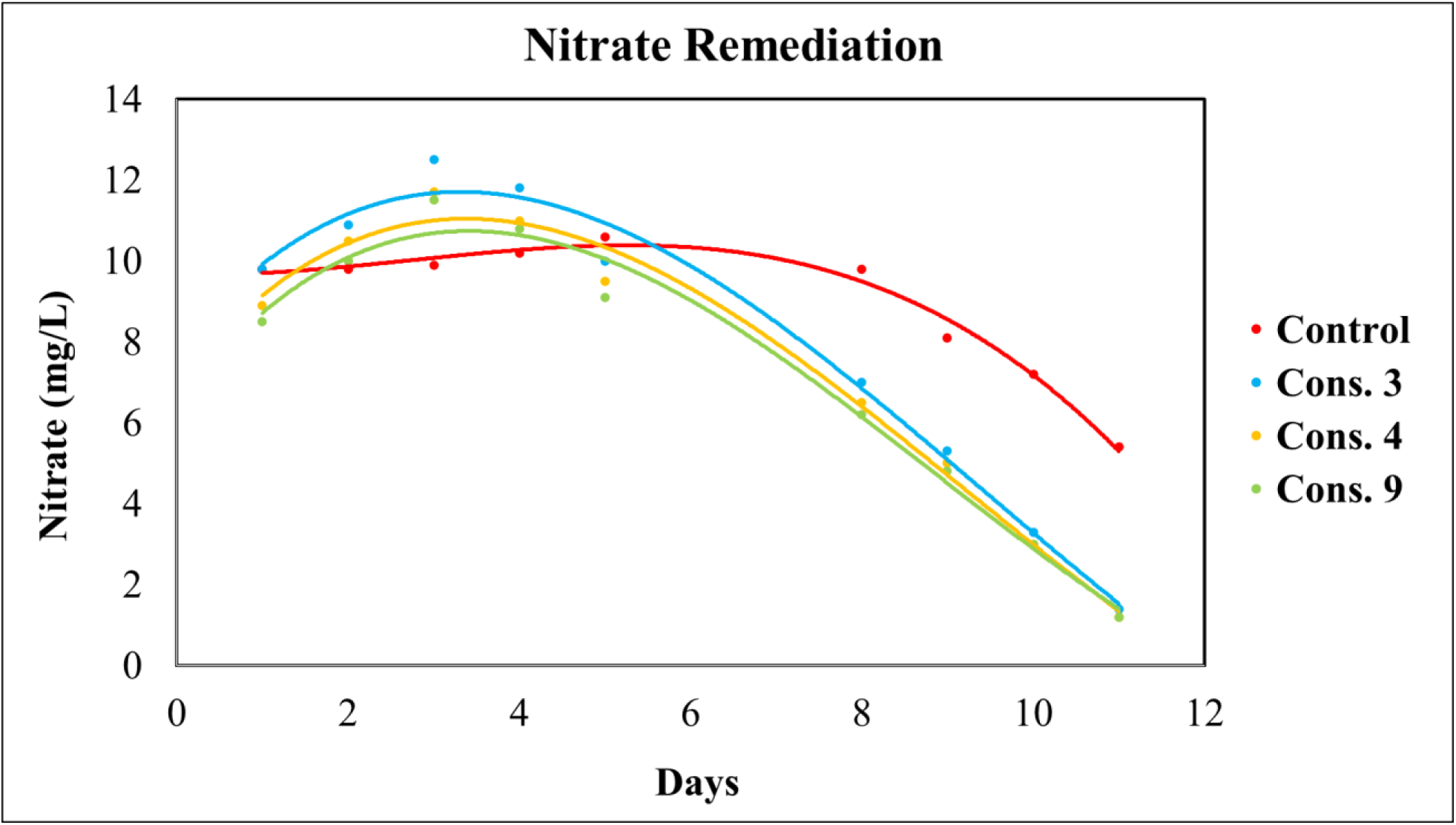
Nitrate removal by the oleaginous consortia isolated from the TP Smith Wastewater Reclamation Facility, Tallahassee, FL. Nitrate content before and after treatment was analyzed using the UV method at 220 and 275 nm with KNO_3_ standard calibration. A control treatment was included with only unautoclaved wastewater sample without addition of isolated consortial strains.

**Figure 9.**
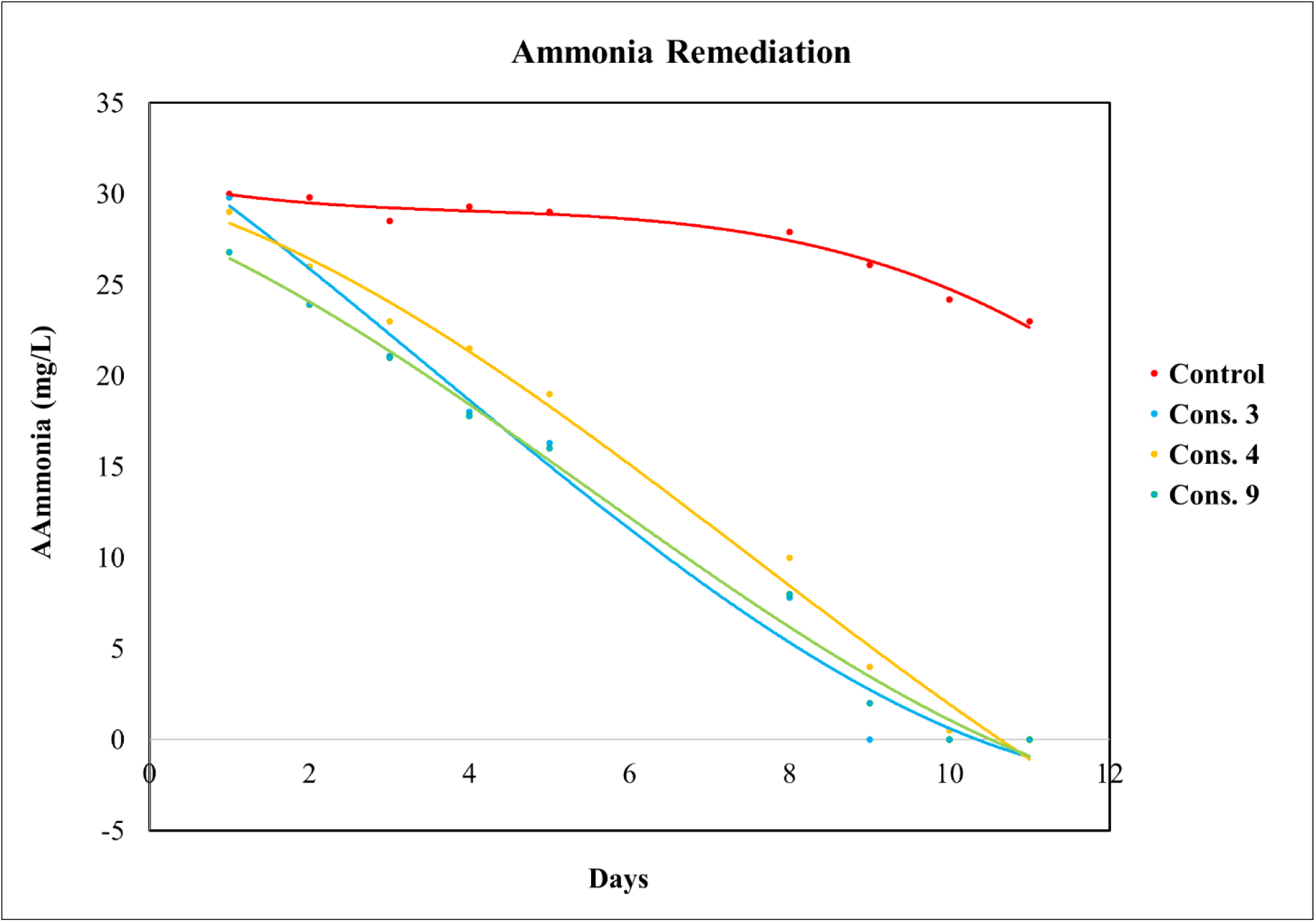
Ammonia removal by the oleaginous consortia isolated from the TP Smith Wastewater Reclamation Facility, Tallahassee, FL. **A**mmonia content before and after treatment was analyzed using a handheld photometer (Lumiso Ammonia Photometer, https://www.waterindustryjournal.co.uk/). A control treatment was included with only unautoclaved wastewater sample without addition of isolated consortial strains.

The nitrogen (N) depletion experiment revealed differential efficiencies among microbial consortia in removing nitrogen from wastewater (Fig. 8). Consortium 9 exhibited the highest nitrogen utilization, characterized by a rapid and substantial decline in nitrogen concentration, indicating robust metabolic activity and effective assimilation or transformation of nitrogenous compounds. Consortium 4 demonstrated moderate performance, with a consistent but less pronounced reduction in nitrogen levels, suggesting functional nitrogen uptake mechanisms with potentially lower metabolic rates or substrate affinity. In contrast, Consortium 3 showed the least efficiency, with minimal nitrogen removal observed throughout the experimental period. The control treatment exhibited negligible changes in nitrogen concentration, confirming that the observed depletion was attributable to microbial activity. These findings highlight Consortium 9 as a promising candidate for nitrogen bioremediation and wastewater treatment applications, while further optimization may be required to enhance the performance of Consortia 3 and 4.

As shown for P and N, the ammonia depletion profile indicates differential capabilities among microbial consortia in removing ammonia from wastewater (Fig. 9). Again, Consortium 9 demonstrated the highest efficiency, with a rapid and pronounced decrease in ammonia concentration, suggesting strong nitrification or assimilation activity. Consortium 4 showed moderate performance, characterized by a steady but less aggressive reduction in ammonia levels, indicating functional but potentially slower metabolic pathways. Consortium 3 exhibited the lowest ammonia removal efficiency, with minimal depletion observed over the experimental period. The control treatment showed negligible change in ammonia concentration, confirming that the observed reductions were attributable to microbial activity. These results position Consortium 9 as a promising candidate for ammonia bioremediation and wastewater treatment, while further optimization may be necessary to enhance the performance of Consortia 3 and 4.

It is noteworthy to mention that the US Environmental Protection Agency generally recommends a limit of 1.0 mg/l total phosphorus and total nitrogen for wastewater effluents. However, limits to nutrient contents of wastewater are being placed and enforced by the state environmental agency depending on the wastewater treatment facility. For example, the T.P. Waste Reclamation Facility (WRF) has a total nitrogen limit of 3 mg/l and total phosphorus of 1.0 mg/l (Pfeffer et al., 2025). The consortial strains isolated in this project have exhibited the capacity to reduce the nutrient contents of the influent to the T.P. Waste Reclamation Facility below the acceptable limit.

## Author Contributions

BE, DA, AP and AC designed experiments. BE performed the lab experiments. AP performed the bioinformatics. BE, AP, DPS and AC wrote the manuscript.

## Acknowledgement

The Florida Department of Agriculture and Consumer Service (FDACs) Office of Energy supported this work through agreement # 24552.

## Supplementary Figures

**Supplementary Figure SI-1.**
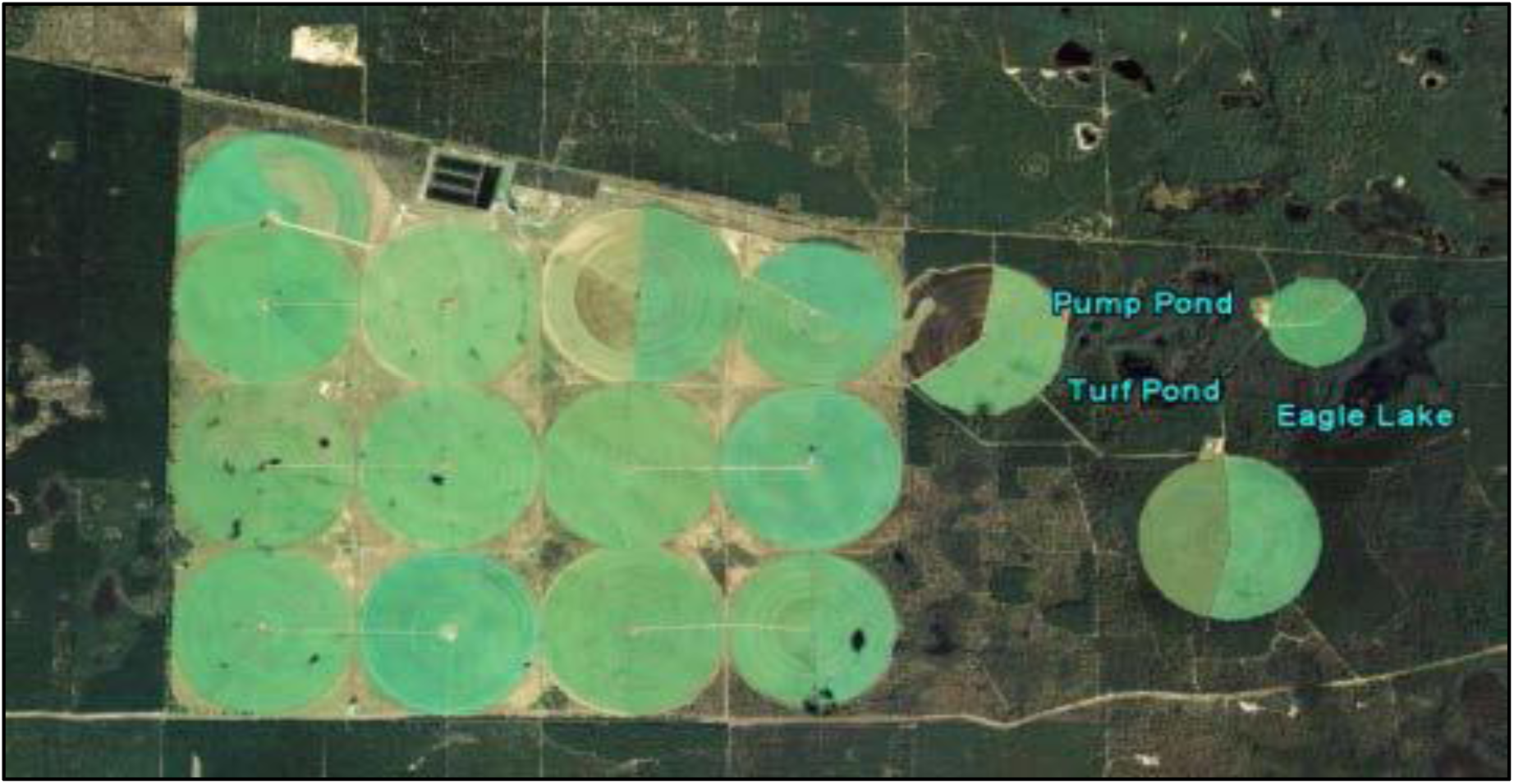
An aerial image of the TP Smith Wastewater Reclamation Facility, Tallahassee, Fl. Three untreated wastewater samples-S1, S2 and S3 for the isolation of oleaginous consortia were collected from the holding tanks and further processed as stated in the main article.

**Supplementary Figure SI-2.**
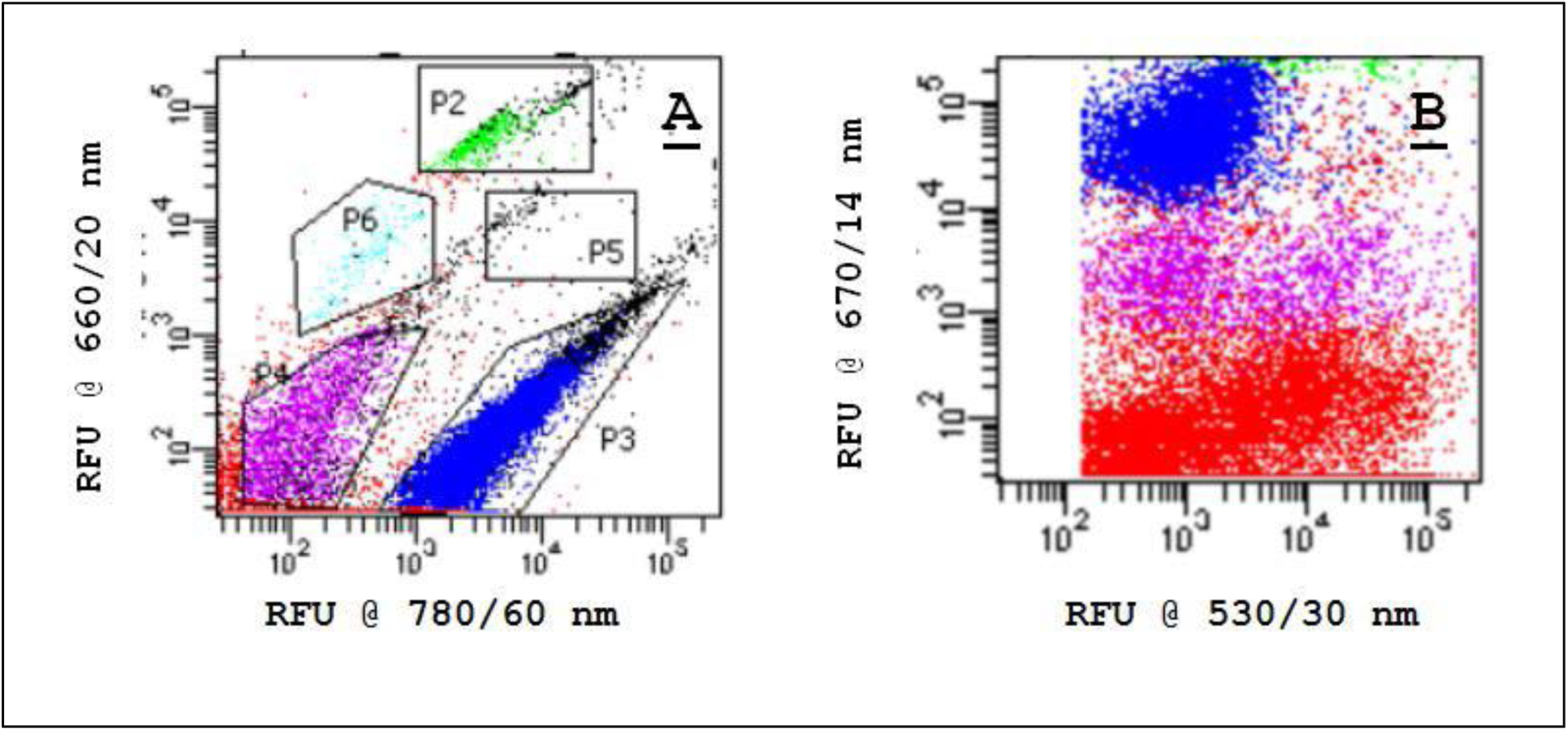
The FACS event log and gating strategy for sample S1. Shown are A, that demonstrates the gating strategy used to select event clusters for the sorting, plotted by the relative fluorescence units (RFU) at 660/20nm against RFU at 780/60nm. The events gated in “A” obtained a unique color set which corresponds to the gate that was carried forward to all plots associated with the sort as shown in B Gate P3 was sorted onto 96-well microtiter plates containing solid BG-11 growth media.

**Supplementary Figure SI-3.**
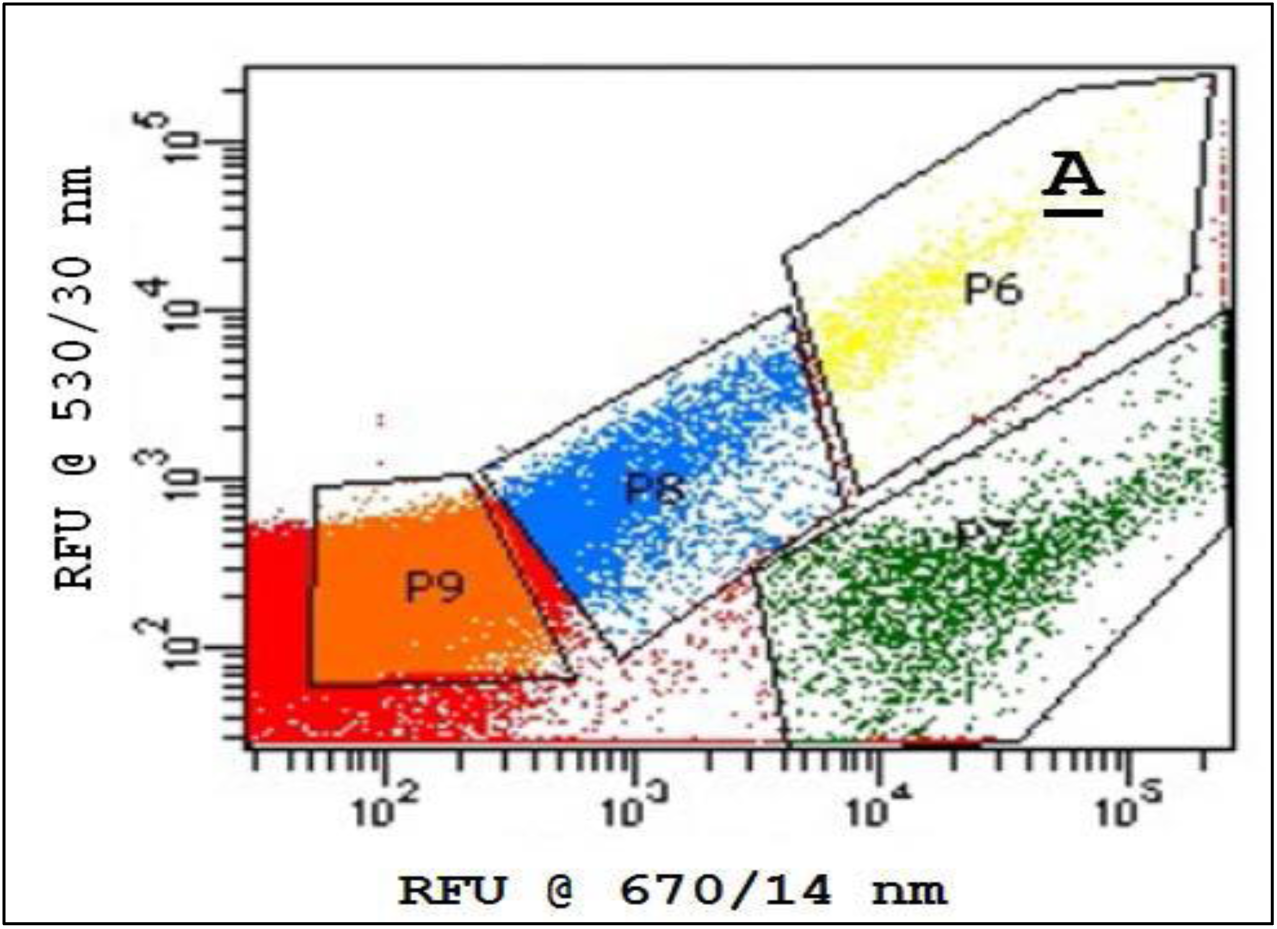
The FACS event log and gating strategy for sample S2. Gates P9, P8, P7, and P6 were created. Gate P6 was sorted at a rate of 1 event per well, onto 96-well microtiter plates containing solid BG11 growth media.

**Supplementary Figure SI-4.**
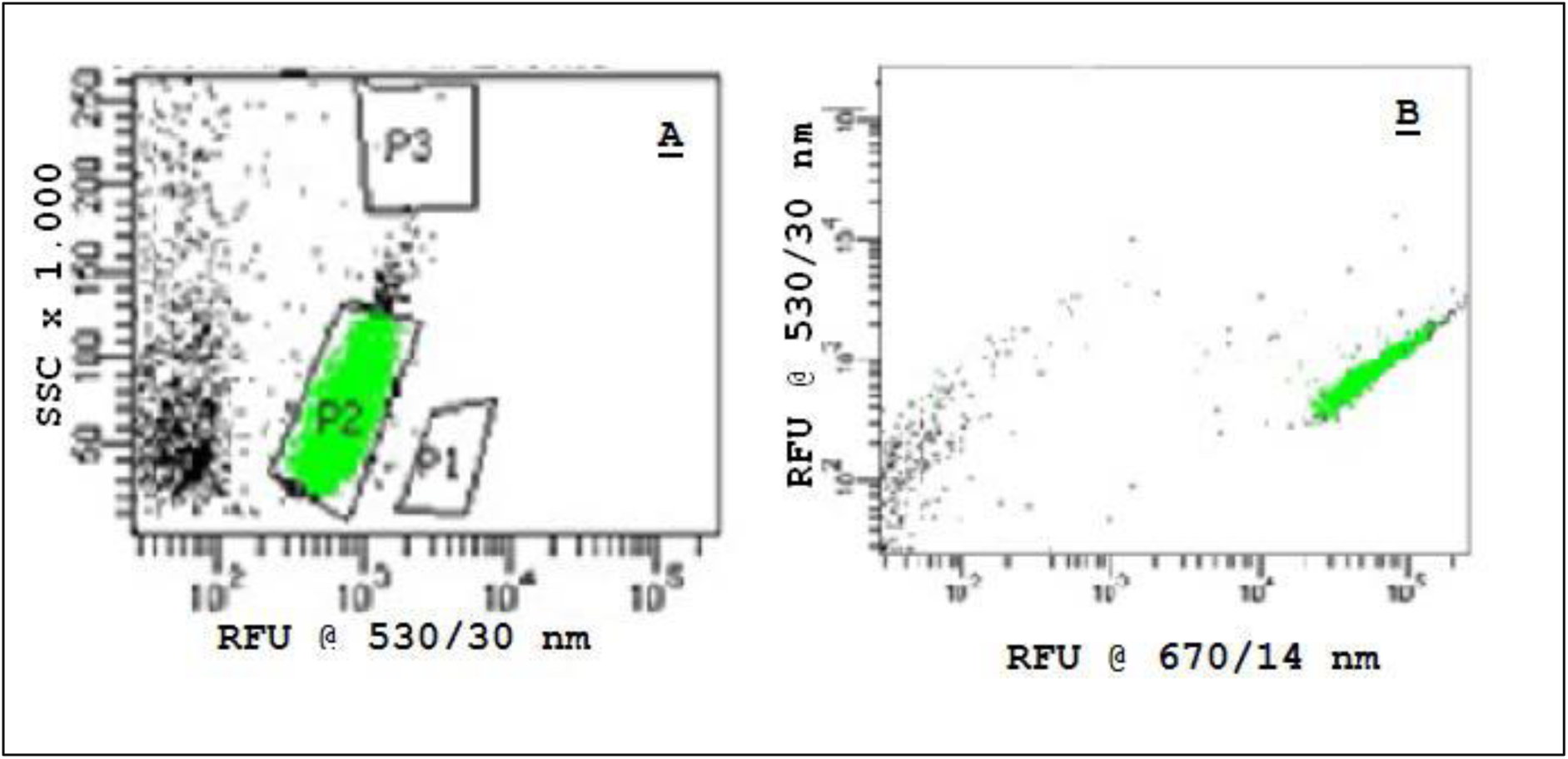
The FACS Event Log and Gating Strategy for sample S3. Graph A plots RFU at 530/30 nm against Side Scatter Complexity (SCC)-a measure of inner cell complexity as a function of light diffraction through the event. The events gated in A obtained a unique color set which corresponds to the gate that was carried forward to all plots associated with the sort as shown in B. Gate P2 was sorted at a rate of 1 event per well onto 96-well microtiter plates containing solid BG11 growth media.

**Supplementary Figure SI-5.**
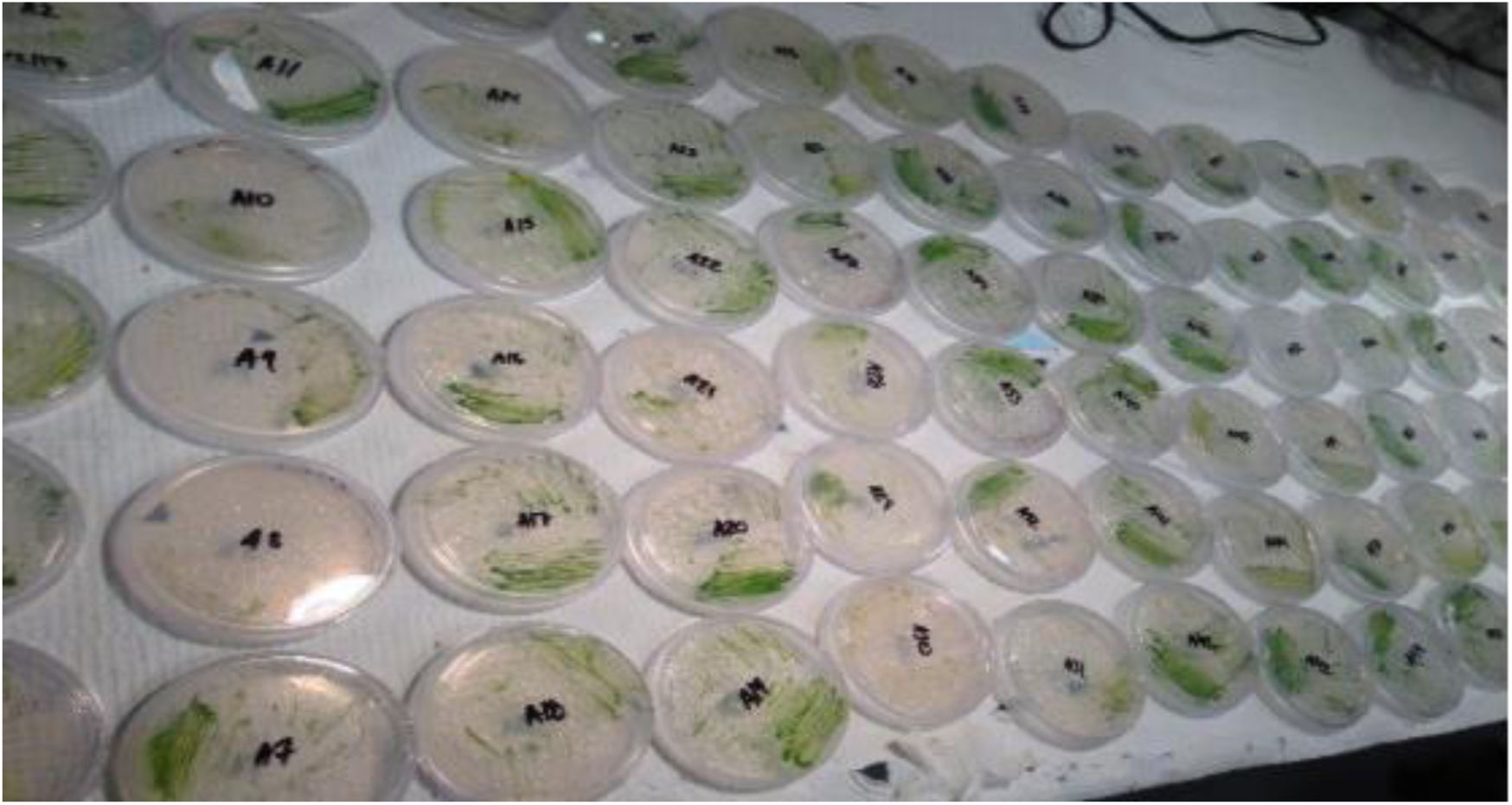
Green colonies that grew on BG11 microtiter plates were further streaked and isolated onto BG11 agar plates for this study.

## References

Acién Fernández, F. G., & María, J. (2018). Recovery of Nutrients From Wastewaters Using Microalgae. Frontiers in Sustainable Food Systems, 2, 396930. 10.3389/fsufs.2018.00059

Andreeva, A., Budenkova, E., Babich, O., Sukhikh, S., Dolganyuk, V., Michaud, P., & Ivanova, S. (2021). Influence of Carbohydrate Additives on the Growth Rate of Microalgae Biomass with an Increased Carbohydrate Content. Marine Drugs, 19(7), 381. 10.3390/md19070381

Antwi, P., Li, J., Boadi, P. O., Meng, J., Koblah Quashie, F., Wang, X., Ren, N., & Buelna, G. (2017). Efficiency of an upflow anaerobic sludge blanket reactor treating potato starch processing wastewater and related process kinetics, functional microbial community and sludge morphology. Bioresource Technology, 239, 105–116. 10.1016/j.biortech.2017.04.124

Assress, H.A., Selvarajan, R., Nyoni, H., Ntushelo, K., Mamba, B.B., Msagati, T.A.M. (2019). Diversity, Co-occurrence and Implications of Fungal Communities in Wastewater Treatment Plants. Sci Rep 9, 14056. 10.1038/s41598-019-50624-z

Berthold, D. E., Shetty, K. G., Jayachandran, K., Laughinghouse, H. D., & Gantar, M. (2019). Enhancing algal biomass and lipid production through bacterial co-culture. Biomass and Bioenergy, 122, 280–289. 10.1016/j.biombioe.2019.01.033

Buchfink, B., Xie, C., and Huson, D. H. (2015). Fast and sensitive protein alignment using DIAMOND. Nat. Methods 12, 59–6010.1038/nmeth.3176.

Cabanelas, I. T. D., Slegers, P. M., Böpple, H., Kleinegris, D. M., Wijffels, R. H., & Barbosa, M. J. (2017). Outdoor performance of Chlorococcum littorale at different locations. Algal Research, 27, 55–64. 10.1016/j.algal.2017.08.010

Cortés-Lorenzo, C., Sipkema, D., Rodríguez-Díaz, M., Fuentes, S., Juárez-Jiménez, B., Rodelas, B., Smidt, H., & González-López, J. (2014). Microbial community dynamics in a submerged fixed bed bioreactor during biological treatment of saline urban wastewater. Ecological Engineering, 71, 126–132. 10.1016/j.ecoleng.2014.07.025

Costa, S., Ganzerli, S., Rugiero, I., Pellizzari, S., Pedrini, P., Tamburini, E. (2017). Potential of Rhodobacter capsulatus Grown in Anaerobic-Light or Aerobic-Dark Conditions as Bioremediation Agent for Biological Wastewater Treatments. Water 9, 108. 10.3390/w9020108

Dao, G., Wu, G., Wang, X., Zhang, T., Zhan, X., & Hu, H. (2018). Enhanced microalgae growth through stimulated secretion of indole acetic acid by symbiotic bacteria. Algal Research, 33, 345–351. 10.1016/j.algal.2018.06.006

Deltac, A. & Meuser, M. (2017). Thomas P. Smith Water Reclamation Facility-Executive Summary. Retrieved March 2025 from https://toxicrisk.com/reports/

Dhariwal A, Chong J, Habib S, King IL, Agellon LB, Xia J.(2017). MicrobiomeAnalyst: a web-based tool for comprehensive statistical, visual and meta-analysis of microbiome data. Nucleic Acids Res. 2017 Jul 3;45(W1):W180–W188. 10.1093/nar/gkx295

Faried, M., Khalifa, A., Samer, M., Attia, Y. A., Moselhy, M. A., Yousef, R. S., Abdelbary, K., & Abdelsalam, E. M. (2023). Biostimulation of green microalgae Chlorella sorokiniana using nanoparticles of MgO, Ca10(PO4)6(OH)2, and ZnO for increasing biodiesel production. Scientific Reports, 13(1), 1–13. 10.1038/s41598-023-46790-w

Gao, Y., Bernard, O., Fanesi, A., Perré, P., & Lopes, F. (2024). The effect of light intensity on microalgae biofilm structures and physiology under continuous illumination. Scientific Reports, 14(1), 1–11. 10.1038/s41598-023-50432-6

Goldstein, R.E. (2015). Green Algae as Model Organisms for Biological Fluid Dynamics. Annual Review of Fluid Mechanics 47, 343–375. 10.1146/annurev-fluid-010313-141426

Gopalakrishnan, K., Wager, Y. Z., & Roostaei, J. (2024). Co-cultivation of microalgae and bacteria for optimal bioenergy feedstock production in wastewater by using response surface methodology. Scientific Reports, 14(1), 1–12. 10.1038/s41598-024-70033-1

Govender, T., Ramanna, L., Rawat, I., & Bux, F. (2012). BODIPY staining, an alternative to the Nile Red fluorescence method for the evaluation of intracellular lipids in microalgae. Bioresource Technology, 114, 507–511. 10.1016/j.biortech.2012.03.024

Gray ND, Miskin IP, Kornilova O, Curtis TP, Head IM. (2002). Occurrence and activity of Archaea in aerated activated sludge wastewater treatment plants. Environ Microbiol. 2002 Mar;4(3):158–68. 10.1046/j.1462-2920.2002.00280.x

Huson, D. H., Auch, A. F., Qi, J., and Schuster, S. C. (2007). MEGAN analysis of metagenomic data. Genome Res. 17, 377–386. 10.1101/gr.5969107.

Huy, M., Kumar, G., Kim, H., & Kim, S. (2018). Photoautotrophic cultivation of mixed microalgae consortia using various organic waste streams towards remediation and resource recovery. Bioresource Technology, 247, 576–581. 10.1016/j.biortech.2017.09.108

Jabari, L., Gannoun, H., Khelifi, E., Cayol, J., Godon, J., Hamdi, M., & Fardeau, M. (2016). Bacterial ecology of abattoir wastewater treated by an anaerobic digestor. Brazilian Journal of Microbiology, 47(1), 73–84. 10.1016/j.bjm.2015.11.029

Karlsson, M., Karlberg, B., & Olsson, R.J. (1995). Determination of nitrate in municipal waste water by UV spectrophotometer. Analytica Chimica Acta, 312 91), 107–113. 10.1016/0003-2670(95)00179-4

Karray F, Ben Abdallah M, Baccar N, Zaghden H, Sayadi S. (2021). Production of Poly(3-Hydroxybutyrate) by Haloarcula, Halorubrum, and Natrinema Haloarchaeal Genera Using Starch as a Carbon Source. Archaea. 2021 Jan 26;2021:8888712. 10.1155/2021/8888712.

Kong QX, Li L, Martinez B, Chen P, Ruan R. (2010). Culture of microalgae *Chlamydomonas reinhardtii* in wastewater for biomass feedstock production. Appl Biochem Biotechnol. 2010 Jan;160(1):9–18. 10.1007/s12010-009-8670-4

Koutra, E., Grammatikopoulos, G., & Kornaros, M. (2017). Microalgal post-treatment of anaerobically digested agro-industrial wastes for nutrient removal and lipids production. Bioresource Technology, 224, 473–480. 10.1016/j.biortech.2016.11.022

Kozyatnyk, I., Benavente, V., Weidemann, E., & Jansson, S. (2025). Adsorption of organic contaminants of emerging concern using microalgae-derived hydrochars. Scientific Reports, 15(1), 1–12. 10.1038/s41598-025-92717-y

Laezza, C., Salbitani, G., & Carfagna, S. (2022). Fungal Contamination in Microalgal Cultivation: Biological and Biotechnological Aspects of Fungi-Microalgae Interaction. Journal of Fungi, 8(10), 1099. 10.3390/jof8101099

Lee, H.-G., Ko, S.-R., Lee, J.-W., Lee, C.S., Ahn, C.-Y., Oh, H.-M., Jin, L. (2017). Blastomonas fulva sp. nov., aerobic photosynthetic bacteria isolated from a Microcystis culture. International Journal of Systematic and Evolutionary Microbiology 67, 3071–3076. 10.1099/ijsem.0.002084

Ma, Q., Qu, Y.-Y., Zhang, X.-W., Shen, W.-L., Liu, Z.-Y., Wang, J.-W., Zhang, Z.-J., Zhou, J.-T. (2015). Identification of the microbial community composition and structure of coal-mine wastewater treatment plants. Microbiological Research, Special Issue: Biodiversity 175, 1–5. 10.1016/j.micres.2014.12.013

Marques, R., Santos, J., Nguyen, H., Carvalho, G., Noronha, J., Nielsen, P. H., Reis, M. A., & Oehmen, A. (2017). Metabolism and ecological niche of Tetrasphaera and Ca. Accumulibacter in enhanced biological phosphorus removal. Water Research, 122, 159–171. 10.1016/j.watres.2017.04.072

Meerbergen, K., Geel, M. V., Waud, M., Willems, K. A., Dewil, R., Impe, J. V., Appels, L., & Lievens, B. (2017). Assessing the composition of microbial communities in textile wastewater treatment plants in comparison with municipal wastewater treatment plants. MicrobiologyOpen, 6(1), e00413. 10.1002/mbo3.413

Mishra, S.K., Suh, W.I., Farooq, W., Moon, M., Shrivastav, A., Park, M.S., Yang, J.-W. (2014). Rapid quantification of microalgal lipids in aqueous medium by a simpl e colorimetric method. Bioresource Technology 155, 330–333. 10.1016/j.biortech.2013.12.077

Neste Corporation (2016). 4 Reasons why the World Needs Biofuels. Retrieved March 2025 from https://www.neste.com/news/4-reasons-why-the-world-needs-biofuels

Numberger, D., Ganzert, L., Zoccarato, L., Mühldorfer, K., Sauer, S., Grossart, H.-P., Greenwood, A.D. (2019). Characterization of bacterial communities in wastewater with enhanced taxonomic resolution by full-length 16S rRNA sequencing. Sci Rep 9, 9673. 10.1038/s41598-019-46015-z

Okoro, V., Azimov, U., Munoz, J., Hernandez, H. H., & Phan, A. N. (2019). Microalgae cultivation and harvesting: Growth performance and use of flocculants - A review. Renewable and Sustainable Energy Reviews, 115, 109364. 10.1016/j.rser.2019.109364

Oswald, W. and Gotaas, H. (1957) Photosynthesis in Sewage Treatment. Transactions of the American Society of Civil Engineers, 122, 73–105.

Pastore M, Sforza E. (2018). Exploiting symbiotic interactions between Chlorella protothecoides and Brevundimonas diminuta for an efficient single-step urban wastewater treatment. Water Sci Technol. 2018 Aug;78(1-2):216–224. 10.2166/wst.2018.155

Pereira, H., Barreira, L., Mozes, A. et al. ((2011). Microplate-based high throughput screening procedure for the isolation of lipid-rich marine microalgae. Biotechnol Biofuels 4, 61. 10.1186/1754-6834-4-61

Pfeffer, K. (2025). Tallahassee WRF: Stringent Limits and Information Modeling. Retrieved March 2025 from https://www.hazenandsawyer.com/projects/process-and-information-modeling-tallahassee-wrf

Pinelli, D., Bovina, S., Rubertelli, G., Martinelli, A., Guida, S., Soares, A., Frascari, D., 2022. Regeneration and modelling of a phosphorous removal and recovery hybrid ion exchange resin after long term operation with municipal wastewater. Chemosphere 286, 131581. 10.1016/j.chemosphere.2021.131581

Plouviez, M., & Brown, N. (2024). Polyphosphate accumulation in microalgae and cyanobacteria: Recent advances and opportunities for phosphorus upcycling. Current Opinion in Biotechnology, 90, 103207. 10.1016/j.copbio.2024.103207

Ramanan, R., Kim, B., Cho, D., Oh, H., & Kim, H. (2016). Algae–bacteria interactions: Evolution, ecology and emerging applications. Biotechnology Advances, 34(1), 14–29. 10.1016/j.biotechadv.2015.12.003

Rayati, M., Rajabi Islami, H., Shamsaie Mehrgan, M. (2020). Light Intensity Improves Growth, Lipid Productivity, and Fatty Acid Pr ofile of Chlorococcum oleofaciens (Chlorophyceae) for Biodiesel Produc tion. BioEnergy Research 13, 1235–1245. 10.1007/s12155-020-10144-5

Ren, H., Xiao, R., Kong, F., Zhao, L., Xing, D., Ma, J., Ren, N., & Liu, B. (2019). Enhanced biomass and lipid accumulation of mixotrophic microalgae by using low-strength ultrasonic stimulation. Bioresource Technology, 272, 606–610. 10.1016/j.biortech.2018.10.058

Rodriguez, D., Santos, K. N., Nagpala, R. T., & Opulencia, R. B. (2023). Metataxonomic Characterization of Enriched Consortia Derived from Oil Spill-Contaminated Sites in Guimaras, Philippines, Reveals Major Role of Klebsiella sp. In Hydrocarbon Degradation. International Journal of Microbiology, 2023, 3247448. 10.1155/2023/3247448

Salwan, R., & Sharma, V. (Eds.). (2020). Physiological and Biotechnological Aspects of Extremophiles. Academic Press

Shaker S, Nemati A, Montazeri-Najafabady N, Mobasher MA, Morowvat MH, Ghasemi Y. (2015). Treating Urban Wastewater: Nutrient Removal by Using Immobilized Green Algae in Batch Cultures. Int J Phytoremediation. 2015;17(12):1177–82. 10.1080/15226514.2015.1045130.

Shivakumar, S., Serlini, N., Esteves, S. M., Miros, S., & Halim, R. (2024). Cell Walls of Lipid-Rich Microalgae: A Comprehensive Review on Characterisation, Ultrastructure, and Enzymatic Disruption. Fermentation, 10(12), 608. 10.3390/fermentation10120608

Silva, G.G.Z., Green, K.T., Dutilh, B.E., Edwards, R.A. (2016). SUPER-FOCUS: a tool for agile functional analysis of shotgun metagenom ic data. Bioinformatics (Oxford, England) 32, 354–361. 10.1093/bioinformatics/btv584

Souza da Silva, C., Pires Santos, G. M., Conceição, G. R., Da Silva Andrade, I., Silva, A. N., Pires Santos, R. M., De Almeida, P. F., & Chinalia, F. A. (2025). Role of low-level alternating current and impedance for enhancing microalgae biomass and lipid production. Journal of Bioscience and Bioengineering. 10.1016/j.jbiosc.2025.02.001

Su, M., Dell’Orto, M., D’Imporzano, G., Bani, A., Dumbrell, A. J., & Adani, F. (2022). The structure and diversity of microalgae-microbial consortia isolated from various local organic wastes. Bioresource Technology, 347, 126416. 10.1016/j.biortech.2021.126416

Tabatabaei, M., Rahim, R. A., Abdullah, N., Wright, A. G., Shirai, Y., Sakai, K., Sulaiman, A., & Hassan, M. A. (2010). Importance of the methanogenic archaea populations in anaerobic wastewater treatments. Process Biochemistry, 45(8), 1214–1225. 10.1016/j.procbio.2010.05.017

Torregrosa-Crespo, J., Martínez-Espinosa, R., Esclapez, J., Bautista, V., Pire, C., Camacho, M., Richardson, D., & Bonete, M. (2016). Anaerobic Metabolism in Haloferax Genus: Denitrification as Case of Study. Advances in Microbial Physiology, 68, 41–85. 10.1016/bs.ampbs.2016.02.001

US Department of Energy (DOE). Biofuel Basics. Retrieved March 2025 from https://www.energy.gov/eere/bioenergy/biofuel-basics

Wang, M., Ye, X., Bi, H. et al. (2024). Microalgae biofuels: illuminating the path to a sustainable future amidst challenges and opportunities. Biotechnol Biofuels 17, 10. 10.1186/s13068-024-02461-0

Wang, M., Zeng, F., Chen, S., Wehrmann, L.M., Waugh, S., Brownawell, B.J., Gobler, C.J., Mao, X., 2024. Phosphorus attenuation and mobilization in sand filters treating onsite wastewater. Chemosphere 364, 143042. 10.1016/j.chemosphere.2024.143042

Wright H. (2022). Phosphorus in wastewater: what is it & why must it be removed? Retrieved from https://www.garrisonminerals.com/post/phosphorus-in-wastewater.

Williams, P. (2007). Biofuel: microalgae cut the social and ecological costs. Nature 450, 478. 10.1038/450478a

Xu, Y., Milledge, J.J., Abubakar, A. et al (2015). Effects of centrifugal stress on cell disruption and glycerol leakage from *Dunaliella salina*. Microalgae Biotechnology, 1(1).pp. 20–27. 10.1515/micbi-2015-0003

Yamane K, Matsuyama S, Igarashi K, Utsumi M, Shiraiwa Y, Kuwabara T. (2013). Anaerobic coculture of microalgae with Thermosipho globiformans and Methanocaldococcus jannaschii at 68°C enhances generation of n-alkane-rich biofuels after pyrolysis. Appl Environ Microbiol. 2013 Feb;79(3):924–30. 10.1128/AEM.01685-12

Zhang, H., Shangguan, M., Zhou, C., Peng, Z., & An, Z. (2023). Construction of a mycelium sphere using a Fusarium strain isolate and Chlorella sp. For polyacrylamide biodegradation and inorganic carbon fixation. Frontiers in Microbiology, 14, 1270658. 10.3389/fmicb.2023.1270658

Zheng, Y., Wan, Y., Zhang, Y., Huang, J., Yang, Y., W Tsang, D. C., Wang, H., Chen, H., & Gao, B. (2022). Recovery of phosphorus from wastewater: A review based on current phosphorous removal technologies. Critical Reviews in Environmental Science and Technology, 53(11), 1148. 10.1080/10643389.2022.2128194

